# Artificial sediments enable reproducible cultivation and recapitulate ecological interactions of cable bacteria

**DOI:** 10.64898/2026.04.15.718496

**Authors:** Kartik Aiyer, Yuang Guo, Lea Emilie Plum-Jensen, Theresa Hagen Van, Marta Lidia Sudo, Robin Bonné, Marko Chavez, Mohamed Y. El-Naggar, Lars Peter Nielsen, Ian P.G. Marshall, Andreas Schramm, Joshua T. Atkinson

## Abstract

Cable bacteria are filamentous microbes that couple sulfide oxidation to oxygen reduction over centimeter distances via long-distance electron transport. While their activity creates characteristic biogeochemical gradients that shape sediment ecology, the study of cable bacteria has been constrained by the chemical and physical heterogeneity of the natural sediments they inhabit. To date, laboratory cultivation efforts have relied on these undefined environmental matrices. Here, we established a reproducible enrichment and cultivation platform using an artificial sediment matrix coupled with chemically defined media. This matrix successfully supported the growth of both freshwater and marine cable bacteria and enabled serial propagation over multiple transfers. Microsensor profiling confirmed that the incubations recapitulated hallmark geochemical signatures, including the sulfide, oxygen and pH gradients associated with electrogenic sulfur oxidation. Scanning electron microscopy confirmed the presence of cable bacteria, while 16S rRNA sequencing confirmed enrichment of the cable bacteria together with a stable co-enriched community that included taxa associated with sulfur and iron cycling as well as cellulose decomposition. This defined cultivation system eliminates the variability inherent to natural samples, providing a controlled platform for dissecting the physiology, genetics, and microbial interactions of cable bacteria.

## Introduction

Cable bacteria are filamentous sulfide-oxidizing bacteria belonging to the family *Desulfobulbaceae* capable of long-distance electron transport (LDET)^1^. They use conductive periplasmic fibers to transport electrons from sulfide oxidizing cells at one end to oxygen reducing cells at the other end. This spatial segregation of redox half reactions over centimeter distances provides competitive advantages to cable bacteria over other sulfide-oxidizers in aquatic sediments and creates characteristic biogeochemical gradients of oxygen, sulfide and pH^2,3^.

Cable bacteria have been discovered to be commonly present in freshwater and marine sediments. Their habitats are diverse, ranging from rivers, lakes, beaches, mangroves^4^, seagrass environments^5^, salt marshes^6^, rhizospheres^7^, groundwater sediments^8^ and continental margins^9^. It was demonstrated that cable bacteria could help in decontamination of petroleum hydrocarbons and could potentially be used in cleaning oil spills^10^. In recent years, they were also shown to reduce methane emissions from rice soil pots, which opens up further possibilities to investigate their ecological roles in mitigating greenhouse gas emissions^11^. As with the majority of environmental microorganisms, the factors that control their growth are poorly understood limiting their ability to be cultivated in defined conditions^12^. Cultivation in defined cultures would facilitate a better understanding of their metabolism as well as enable mass production for environmental engineering applications and potentially for genetic manipulation of cable bacteria.

Currently, the most widespread approach to cultivating cable bacteria in labs is as enrichments in environmentally derived sediments^13,14^. In this method, a single cable bacteria filament from an environmental source is inoculated into partially sterilized natural sediments and allowed to grow along with other microbes. By serially transferring sediment clumps into fresh sterilized sediments, a single-strain enrichment can be established and propagated for lab-based experiments^15^. One challenge with this approach is the spatiotemporal heterogeneity of environmental samples. This results in sediments with varying abiotic (*e.g.,* porosity, salinity, pH, buffering capacity, mineral and organic matter composition) and biotic (*e.g.,* microbial and viral communities) properties. This sample heterogeneity has been a major limitation for the study of cable bacteria making it challenging to reproduce studies and directly link changes in geochemistry to the activity of cable bacteria.

The complex composition of environmental samples makes it challenging to collect large quantities of cable bacterial biomass that are amenable for subsequent physiological studies including proteomics and ultrastructural analysis. Attempts have been made to cultivate cable bacteria in sediment-free gradient agar tubes that are used to cultivate other filamentous sulfur oxidizing bacteria such as *Beggiatoa,* but to-date have been unsuccessful^16,17^. As alternatives, inert matrices, such as glass beads^1,18^ or agar pillars^19^, have been amended to sediments to aid the separation of cable bacteria from sediments particles. These approaches exploit the gliding motility of cable bacteria, allowing them to migrate into the inert matrix away from sediment particles, while still depending on the native environmental sediment for growth^20^. Recently, a bentonite-based medium in a bioreactor was described to grow cable bacteria away from the natural sediment^21^. However, no attempt was made to propagate these cultures or evaluate the metabolic activity of the cultures when grown in the bentonite medium.

Using artificial sediments to cultivate environmental microbes has the potential to reveal the impact of the physicochemical properties of the sediment on their growth. Studies using standardized artificial soils with varying texture, minerology, pH, and organic matter content have revealed impacts on the growth and chemical signaling between microbes using acyl-homoserine lactones for quorum sensing^22^. Similar approaches using various mineral phase and organic compounds have been applied to recapitulate aquatic sediment environments primarily to develop standardized ecotoxicity studies^23–25^. However, artificial sediments have largely been applied to standardize the growth of aquatic plants and benthic fauna and have not been evaluated for cultivating cable bacteria or other microbes that are recalcitrant to conventional cultivation approaches. Developing a reproducible and chemically defined sediment system would provide a controlled experimental platform to investigate cable bacteria physiology and microbial interactions.

Here, we evaluated enrichment and continuous culturing of cable bacteria in artificial sediments immersed in a defined nutrient medium. Two artificial sediment matrices were prepared containing either: (i) insoluble organic components with freshly precipitated iron sulfide or (ii) a mixture of sand, clay, insoluble organic components, and freshly precipitated iron sulfide. These artificial sediments were designed with a matrix for cable bacteria attachment and motility and with a nutrient medium providing the essential components for growth. We measured the biogeochemical profiles of oxygen, sulfide and pH in the artificial sediments, evaluated filament morphology using microscopy, and performed 16S rRNA gene sequencing to identify other members of the community. We successfully cultivated cable bacteria through six serial transfers between artificial sediments. The use of artificial sediments eliminates the heterogeneity constraints imposed by natural sediments and could be useful in studying metabolic capabilities and the ecological interactions of cable bacteria with other microbes as well as providing a system to reproducibly generate cable bacteria biomass.

## Materials and Methods

### Preparation of the nutrient medium

Artificial sediments were washed and incubated in a freshwater benthic medium (FWBM). FWBM has a defined chemical composition and is based on media used to cultivate *Beggiatoa*, another group of filamentous sulfur-oxidizing bacteria^16^. FWBM consisted of the following (per 1000 mL, see **Table S1**): calcium sulfate dihydrate (0.12 g), ethylenediaminetetraacetic (EDTA) (0.01 g), magnesium sulfate heptahydrate (0.2 g), sodium chloride (0.016 g), calcium chloride dihydrate (0.264 g), sodium phosphate dibasic (0.071 g), potassium phosphate monobasic (0.068 g), sodium bicarbonate (0.168 g), sodium acetate (0.004 g), ammonium chloride (0.003 g), sodium nitrate (0.004 g), ferric chloride (2 mL of 0.29 g L^−1^ solution), 1 mL of 1000x micronutrient solution (**Table S2)**, and 0.416 mL of 2400x vitamin solution (**Table S3)**.

### Preparation of FeS slurry

Amorphous FeS was freshly precipitated by dissolving ferrous sulfate heptahydrate (46.2 g) and sodium sulfide nonahydrate (39.6 g) into 300 mL of 50**°**C deionized water that was rapidly stirred^26^. A thick black precipitate forms instantaneously. The mixture is mixed for 3 min to completely dissolve ferrous sulfate and sodium sulfide. The FeS precipitate was decanted into a narrow-mouthed glass bottle (500 mL), filled to the top with oxic deionized water and stoppered tightly. The FeS was allowed to settle for 2 h, following which overlaying water was decanted and replaced five times until the pH of the precipitate was neutral. The final concentration of the well-mixed FeS slurry was ∼0.33 M.

### Preparation of artificial sediment

Two separate artificial sediment recipes, AS1 and AS2, were formulated to test the growth of cable bacteria. AS1 consisted of exclusively alpha cellulose which served as insoluble organic matrix^23^. AS2 consisted of a mixture of alpha cellulose, sand, and kaolinite clay. To prepare AS1, alpha cellulose (0.4 g, Sigma Cat. No. C8002) was weighed into a 50 mL centrifuge tube. To prepare AS2, alpha cellulose (0.4 g, Sigma Cat. No. C8002), 50-70 mesh quartz sand (17.6 g, Sigma Cat. No. 274739), and kaolinite clay (2 g, Sigma Cat. No. 03584) were weighed into a 50 mL centrifuge tube. For both AS1 and AS2, 20 mL of the FWBM and 5 mL of iron sulfide slurry were added and tubes were allowed to equilibrate on a tube roller for 1 h at 25**°**C. To settle the mixture to the bottom of the tubes, tubes were centrifuged for 5 min at 1000 rcf in a swinging-bucket rotor. Both AS were incubated in water bath (90°C) for 60 min. The tubes were subsequently washed with 20ml of FWBM and centrifuged. The pH of the mixture after washing was 7. After washing, the sediment was transferred to 20 mL plastic cups along with the overlay medium.

### Artificial sediment inoculation for freshwater cable bacteria

To initiate primary enrichments, artificial sediments were inoculated with ∼200 mg of freshwater sediment from an enrichment of *Electronema aureum* GS that is maintained at Aarhus University^13^. After inoculation, the artificial sediment tubes were incubated in an aquarium filled with FWBM for *E. aureum* (AS1-F and AS2-F) that was continuously aerated using an aquarium stone and air pump at 18°C. Prior to the addition of the sediments, the aquaria were allowed to equilibrate overnight to reach a stable oxygen concentration. To estimate the suitability of artificial sediments for continuous culturing of cable bacteria, serial transfers of artificial sediments were performed for *E. aureum*, every 14 days by transferring ∼200 mg of artificial sediments containing active cable bacteria to fresh sediments. Artificial sediment enrichments were established in triplicates, while the natural sediment inoculum was analyzed as a single reference sample.

### Oxygen, sulfide and pH microsensor profiling

Depth profiles of pH, O_2_, and H_2_S were obtained immediately after inoculation and 14 days after inoculation using a multichannel Microsensor Multimeter (UniAmp fx-6, Unisense)^2^. Microsensor (pH and H_2_S) or optode probes (O_2_) were mounted on a microprofiling system to facilitate incremental movements to generate vertical profiles. SensorTrace PRO software (Unisense) was employed to manage the microsensor movement and to record the sensor data. Microsensors were calibrated immediately before measurements.

### Trench-slide microscopy

To visualize cable bacteria in the artificial sediments via microscopy, custom-made trench slides were used. Trench slides were generated by sand-blasting a 1 mm deep trench into a 2 mm thick glass slide^15,20^. 14 days after inoculation, trench slides were prepared by transferring a portion of the upper 2 cm^3^ from artificial sediment into the trench. 0.5 mL of anaerobic FWBM was placed on top of the sediment and a cover slip was floated over top and sealed using polydimethylsiloxane (PDMS, Sylgard 184 7:1). To allow for the formation of an oxygen gradient and migration of cable bacteria out of the sediment, the trench slides were incubated for 24-48 h and then were observed under a microscope using dark field.

### Scanning electron microscopy (SEM)

SEM was performed on cable bacteria extracted from artificial sediments using a dual-beam FIB/SEM 2D Versa (ThermoFisher). Cable bacterium filaments were fished out of the cultures with glass hooks and cleaned in 5 consecutive MilliQ droplets. In total 100 to 200 filaments were collected in a 1.5 mL centrifuge tube and directly afterwards pipetted on SiO_2_ wafers for SEM imaging. A vacuum of 5 kV, a working distance of 10mm and a current of 13pA consisted of the acquisition parameters.

### 16S rRNA amplicon sequencing for freshwater cable bacteria

For the *E. aureum* cultures, the microbial community growing in the artificial sediments was analyzed via 16S rRNA sequencing. 0.25 g of the upper 0–6 mm of the sediment was carefully collected using sterile spatulas. After collection, the sample was homogenized prior to DNA extraction. DNA was extracted from 0.25 g of artificial sediments or natural sediments using the DNeasy PowerLyzer PowerSoil kit according to manufacture instructions (Qiagen, Hilden, Germany), and the V3-V4 region of bacterial 16S rRNA genes was amplified using primers Bac 341F and Bac 805R^27^. The 16S rRNA gene amplicon libraries were prepared according to Illumina’s 16S Metagenomic Sequencing Library Preparation guide with three consecutive PCR reactions for amplification of the target regions (1^st^ PCR: 20 cycles), addition of adapters (2^nd^ PCR: 10 cycles) and indexes (3^rd^ PCR: 8 cycles). The amplified DNA was sequenced on a MiSeq bench top sequencer (Illumina) and analyzed with the DADA2 pipeline (v. 1.27.1)^28^ using the SILVA database (version 138.1). Raw sequences can be downloaded from the NCBI/EMBL-EBI/DDBJ Sequence Read Archive under BioProject ID PRJNA1453017.

### Sediment sampling and generation of a clonal enrichment culture of marine cable bacteria

*Candidatus* Electrothix sp. NJ1 was enriched from a marine sediment obtained from a tidal mud flat found at Crabtown Creek in Fisherman’s Cove Conservation Area, Manasquan, NJ, USA (40°06’28.2“N 74°02’41.0“W). The upper 20 cm of sediment was harvested. Sediments were transported to Princeton University and stored at 25°C in sealed containers with overlaying water for 28 h. Sediment was then homogenized and sieved through a 1 mm mesh screen. To reduce the initial sediment biomass the sediment was autoclaved at 121°C for 2 h. Autoclaved sediment was packed into a glass shell vial (Fisherbrand Cat. No. 03-339-26F) and inoculated with a 3 cm^3^ portion of an undisturbed natural sediment from the field that was microscopically inspected for the presence of cable bacteria. To enrich for cable bacteria, the inoculated core was incubated for 3 months in an aerated aquarium containing water from the field site that was filtered using a 0.2 µm PES filter. The overlaying seawater was measured to have a salinity of 25 ppt measured using a refractometer. Single cable bacteria filaments were obtained from this enrichment and used to inoculate fresh autoclaved sediments to establish enrichments that were serially passaged and enriched in autoclaved sediment 3 times prior to use in this study^13^.

### Artificial sediment inoculation for marine cable bacteria

In place of FWBM, an artificial seawater medium (ASM) was generated by dissolving instant ocean (Spectrum Brands, Blacksburg, VA, USA) at 25 ppt salinity supplemented with 1 mL of 1000x micronutrient solution (**Table S2**) of 0.416 mL of 2400x vitamin solution (**Table S3**) per 1000 mL. For marine AS2 sediments, 20 mL of ASM and 5 mL of iron sulfide slurry were added to each tube and equilibrated on a tube roller for 1 hour at 25 °C. To initiate primary enrichments, artificial sediments were packed into 25 mL glass shell vials (Fisherbrand Cat. No. 03-339-26F) and inoculated with ∼200 mg of the enrichment of *Ca*. Electrothrix sp. NJ1 from this study. After inoculation, the artificial sediment (AS2) tubes were incubated in an aquarium filled with ASM that was continuously aerated using an aquarium stone and air pump at 23°C. Prior to the addition of the sediments, the aquaria were allowed to equilibrate overnight to reach a stable oxygen concentration. To estimate the suitability of artificial sediments for continuous culturing of cable bacteria, serial transfers of artificial sediments were performed for *Ca*. Electrothrix sp. NJ1, every 30 days, by transferring ∼200 mg of artificial sediments containing active cable bacteria to fresh sediments.

### 16S rRNA amplicon sequencing for marine cable bacteria

For the *Ca.* Electrothrix sp. NJ1 cultures, the microbial community growing in the natural sediment was analyzed via 16S rRNA sequencing. Samples were collected and immediately frozen at −80 °C overnight. DNA was extracted from approximately 0.4 g of artificial sediment using the DNeasy PowerLyzer PowerSoil Kit (Qiagen, Hilden, Germany), following the manufacturer’s instructions with minor modifications: a phenol:chloroform:isoamyl alcohol extraction (25:24:1, Sigma, Cat. No. 77617) was performed after bead beating, followed by two washes with chloroform:isoamyl alcohol (24:1, Sigma, Cat. No. C0549). Full-length 16S rRNA genes were amplified using universal primers 27F and 1492R^29^ with Q5 High-Fidelity DNA Polymerase (NEB, Cat. No. M0491) under 32 PCR cycles consisting of denaturation at 98 °C for 45 s, annealing at 55 °C for 45 s, and extension at 72 °C for 90 s. Amplicons were sequenced using Oxford Nanopore Technologies using the v14 library prep chemistry, R10.4.1 flow cells, and the super accurate basecalling model through the Plasmidsaurus Microbiome Amplicon service. Raw sequences can be downloaded from the NCBI/EMBL-EBI/DDBJ Sequence Read Archive under BioProject ID PRJNA1453017.

### Phylogenetic analysis

The full-length 16S rRNA gene sequence of *Ca.* Electrothrix sp. NJ1 was recovered from nanopore amplicon data by aligning reads against a curated reference database of 90 cable bacteria 16S rRNA gene sequences (**Appendix A**). High-confidence candidate sequences were identified by trimming to reads between 1200-1600 bp aligning to the cable database using minimap2 (v2.30-r1287)^30^ and selecting matches with a minimum identity threshold of 98%. We aligned these hits to their matches in the cable bacteria database using MUSCLE (v5.3)^31^. The 16S rRNA gene sequence the NCBI/EMBL-EBI/DDBJ GenBank under BioProject ID PRJNA1453017.

For phylogenetic analysis, representative 16S rRNA gene sequences (>1200bp) of 89 cable bacteria and 18 related taxa from Ley et al^32^ as well as the recently identified *Ca*. Electrothrix yaqonensis YB6^33^ and *Ca*. Electrothrix sp. NJ1 from this study were aligned using MUSCLE(v5.3)^31^. A maximum likelihood tree was constructed with IQ-TREE (v1.6.12)^34^, employing the TIM3e+I+G4 model and 1,000 ultrafast bootstrap replicates. The sequence of *Geobacter sulfurreducens* PCA (NR_075009) was used as an outgroup. Resulting phylogenies were visualized using iTOL (v6)^35^. Pairwise percent identities were calculated using the MUSCLE alignment and clustered based on the phylogenetic tree.

## Results and Discussion

### Establishment of artificial sediment enrichments for freshwater cable bacteria

To evaluate if artificial sediments support the growth of cable bacteria, both AS1 and AS2 were inoculated with a 200 mg sample from the upper 2 cm^3^ of a natural sediment containing an enrichment of *E. aureum* GS. The electrogenic sulfur oxidation (e-SOx) activity of cable bacteria was monitored by measuring the vertical chemical profiles of sulfide, oxygen, and pH using microsensor probes both immediately following inoculation and following 14 days of incubation in an aerated aquarium (**Figure 1**). At the initial time point, both AS1 and AS2 displayed similar geochemical profiles with <10 µM H_2_S and a pH of 7.5 that was invariant throughout the upper 6 mm of the sediment core, while the concentration of O_2_ linearly decreased from ∼250 μM at the surface to 0 μM at ∼3 mm depth (**Figure S1**). This indicates that the chemical conditions of the two sediment formulations in the absence of cable bacteria are similar, which is expected as the two formulations only differ in the matrix material.

**Figure 1:**
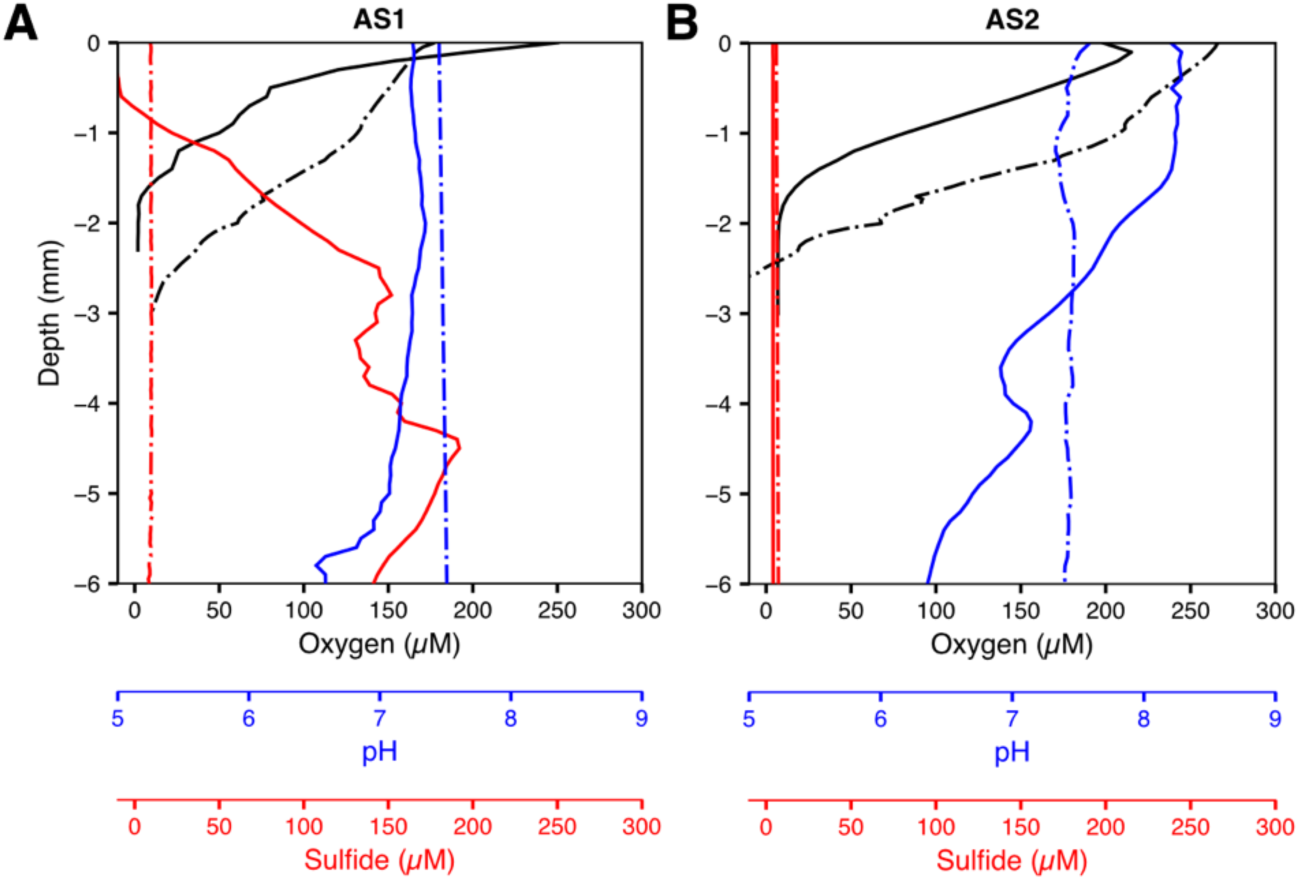
Vertical chemical profiles in artificial sediments with freshwater cable bacteria. Oxygen (black), sulfide (red) and pH (blue) profiles were measured for either **(A)** AS1 or **(B)** AS2 sediments inoculated with enrichments of freshwater *E. aureum* (solid line) or non-inoculated controls (dash-dot line) following 14 days of incubation in aerated FWBM.

Following 14 days of incubation, the oxygen penetration depth and oxic-zone pH of AS1 sediments inoculated with the *E. aureum* enrichment remained similar to uninoculated sediments (**Figure 1**) and the inoculated sediments at the initial time point (**Figure S1**). In AS1, an overlapping region of about 1.5 mm containing free sulfide and dissolved oxygen suggested that LDET and e-SOx metabolism is not occurring or is marginal within this sediment. AS2 sediments displayed pH profiles that were elevated to ∼8 in the oxic zone. The sulfide profiles of AS2 were also markedly different than AS1, for AS2 there was no detectable increase in sulfide throughout the 6 mm of the core, indicating a suboxic zone of at least 4 mm depth. The elevated pH indicating proton consumption in the oxic zone and the formation of a suboxic zone are hallmark geochemical signatures of e-SOx metabolism mediated by cable bacteria^3^. These results suggest that AS2 can support the e-SOx activity of *E. aureum*, while AS1 does not support e-SOx activity. This may be the result of differing sediment porosity resulting from the varying particle sizes of alpha-cellulose (200-500 μm) used in AS1 relative to the sand (200-300 μm) and clay particles (<2 μm) included in AS2 that may impact oxygen diffusion or cable bacteria motility. Additionally, the inclusion of the clay particles may enhance the cation exchange capacity and nutrient adsorption of the sediment matrix.

### Continuous culturing in artificial sediments

To evaluate if artificial sediments can be used for growth and continuous culturing of cable bacteria, we performed serial transfers of sediments from both AS1 and AS2 primary enrichments into fresh artificial sediments. For each serial enrichment, sediments were allowed to develop over 14 days prior to measurement of chemical profiles and transfer to a fresh artificial sediment. The secondary (E2) and tertiary (E3) enrichments in both AS1 and AS2 resulted in geochemical profiles that were similar to primary enrichments (E1) (**Figure 2**). For AS1 sediments, the oxygen and sulfide always overlapped at a depth of ∼1.5-2 mm with no indication of sulfide consumption and the pH was stable throughout the sediment. In contrast, in AS2 sediments, a ∼1.5 mm suboxic zone developed, marked by a pH minimum in the anoxic region near the advancing sulfide front and an elevated pH (∼8) near the oxic surface, reflecting the distinct half-reactions catalyzed by cable bacteria (sulfide oxidation in the anoxic zone and oxygen reduction in the oxic zone). We evaluated if AS2 sediments would support long-term cultivation of cable bacteria by subculturing an additional three times. Because AS1 was not found to support cable bacteria activity or growth continued subcultures of this sediment were not performed. Further subculturing (cultures 4-6) gave similar results (**Figure S2**). These results suggest that the AS2 formulation can support long-term serial growth and replication of cable bacteria, while the AS1 formulation does not support growth.

**Figure 2.**
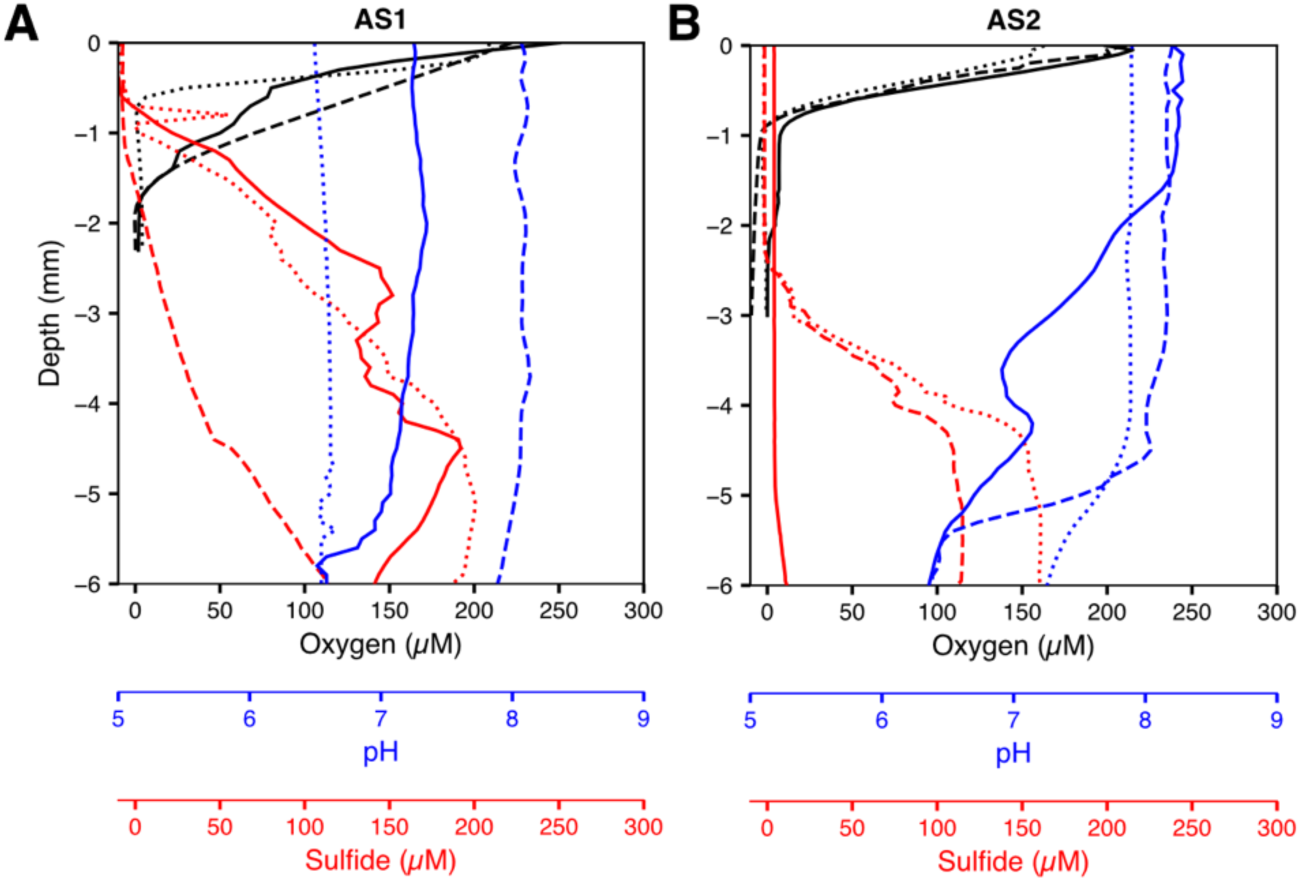
Vertical chemical profiles in artificial sediments for multiple transfers. Oxygen (black), sulfide (red) and pH (blue) profiles were measured for **(A)** AS1 or **(B)** AS2 sediments with serial enrichments of freshwater *E. aureum* following 14 days of incubation in aerated FWBM. For reference, the primary enrichment E1 from Figure 1 is shown (solid line) along with the secondary enrichment E2 (dotted line), and the tertiary enrichment E3 (dashed line).

### Evaluating morphology of cable bacteria grown in artificial sediments

To complement the geochemical profile measurements, we used microscopy to investigate the morphology and behavior of cable bacteria grown in artificial sediments. Cable bacteria were harvested from AS2 enrichments of *E. aureum* and were evaluated using scanning electron microscopy. SEM imaging confirmed that the filamentous bacteria present in AS2 matched the morphology of *E. aureum*^15,36^. AS2 grown filaments were ∼ 1 μm in diameter and had about 40 parallel ridges running the length of the cell that contain the periplasmic conductive fibers (**Figure 3A**). Interconnected cells formed multiple cell septa likely undergoing coordinated cell division in the filament (**Figure 3B**) as has been observed using nanoscale secondary ion mass spectrometry (nanoSIMS)^37^, confirming that cable bacteria grow and actively divide in AS2. A dense network of intertwined cable bacterial filaments displayed uniform morphologies suggesting similar growth throughout the cable bacteria population (**Figure 3C**).

**Figure 3:**
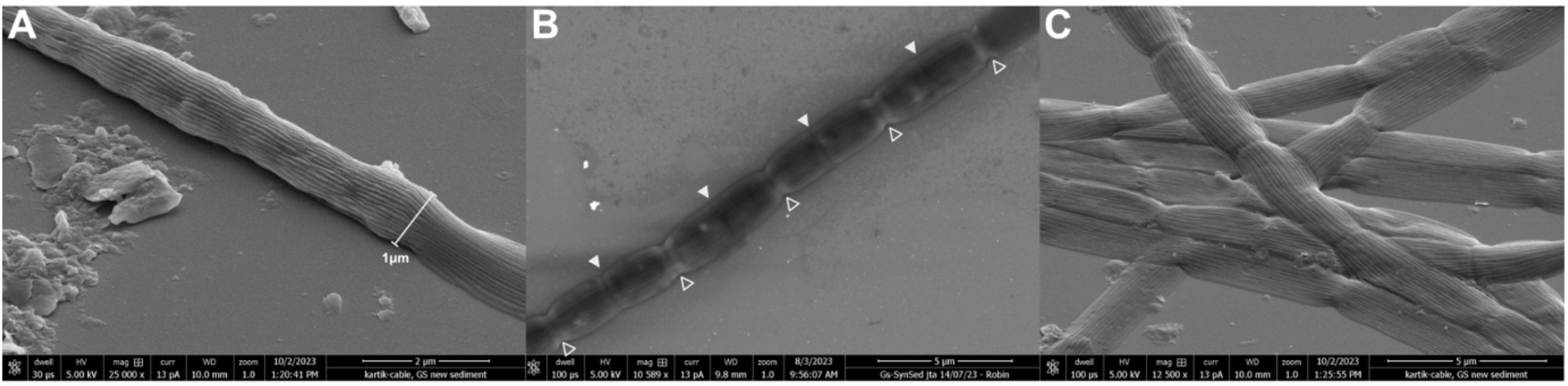
Scanning electron micrographs of *E. aureum* enriched in AS2. **(A)** Filaments show typical surface ridge morphology with a diameter for 1μm. **(B)** Filaments display both new (closed triangles) and old division planes (open triangles). **(C)** Bundles of filaments display uniform morphology.

To evaluate if cable bacteria behave similarly within the artificial sediments as in natural sediments, we performed live cell microscopy on trench slides^15^ prepared with AS2 containing *E. aureum* on their sixth serial enrichment. Cable bacteria were seen emerging from the sediment and aggregating (**Figure 4**), which is typical cable bacteria behavior. Additionally, the phenomenon of single-celled bacteria “flocking” toward active cable bacteria filaments was also observed in artificial sediment enrichments (**Video S1**). This behavior is consistent with earlier observations described in natural sediments^38^. This suggests that cable bacteria-associated microbes have been propagated together with cable bacteria in the artificial sediment, and that ecological interactions such as flocking can be recapitulated in these artificial environments. Collectively the similar morphologies and behaviors indicate that cable bacteria grown in artificial sediment environments are growing consistently as those grown in natural sediments.

**Figure 4:**
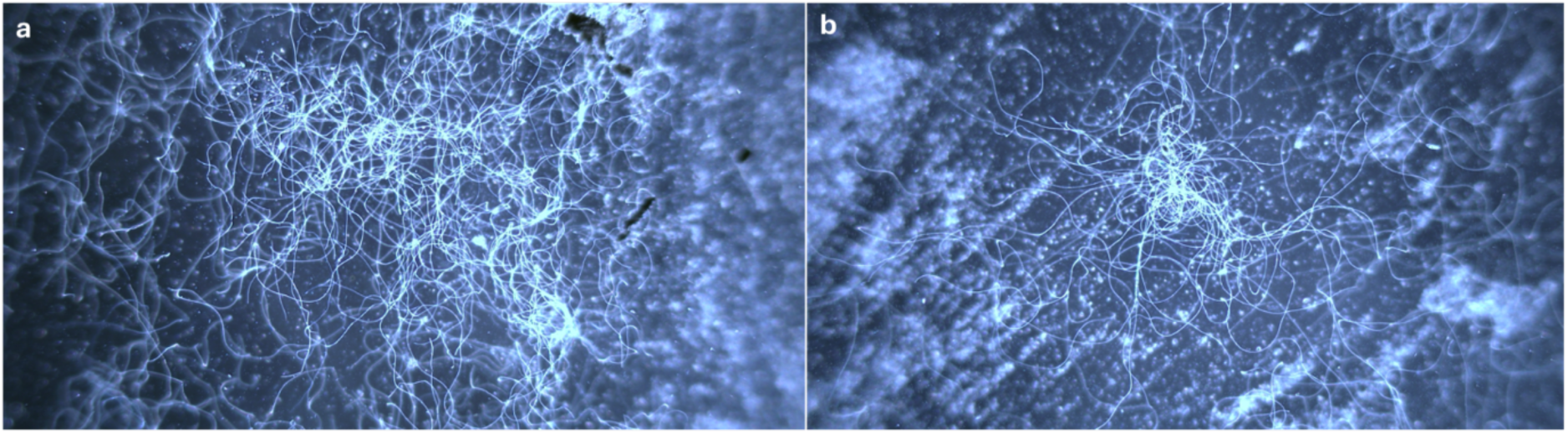
Dark field microscopy of *E. aureum* bundles observed in trench slides.

### Evaluating the microbial community during serial enrichments

To assess how the microbial community composition changed during enrichment in artificial sediments, we harvested DNA from the first three passages in both AS1 and AS2 as well as the natural sediment inoculum and performed partial 16S rRNA gene sequencing. We first evaluated the relative abundance of *E. aureum* across all conditions (**Figure 5**). For AS2, a higher relative abundance of *E. aureum* is observed in the primary (E1) and tertiary (E3) enrichments, while *E. aureum* was detected at a lower relative abundance in the secondary enrichments (E2). The reduction of *E. aureum* reads in E2 is attributed to sampling bias as the geochemical profile (**Figure 2**) and microscopy of sample indicated cable bacteria activity. By contrast, AS1 sediments poorly encouraged the growth of cable bacteria with *E. aureum* reads detected only in the primary enrichment and no reads were detected in the secondary or tertiary enrichments. This corroborates the results from geochemical profiling and microscopy. The first inoculation into artificial sediments consisted of a clump transfer containing an abundance of cable bacteria, which likely explains their abundance in E1 for both AS1 and AS2. In all cases, including the natural sediments, the cable bacteria were less than 10% of the total observed reads.

**Figure 5.**
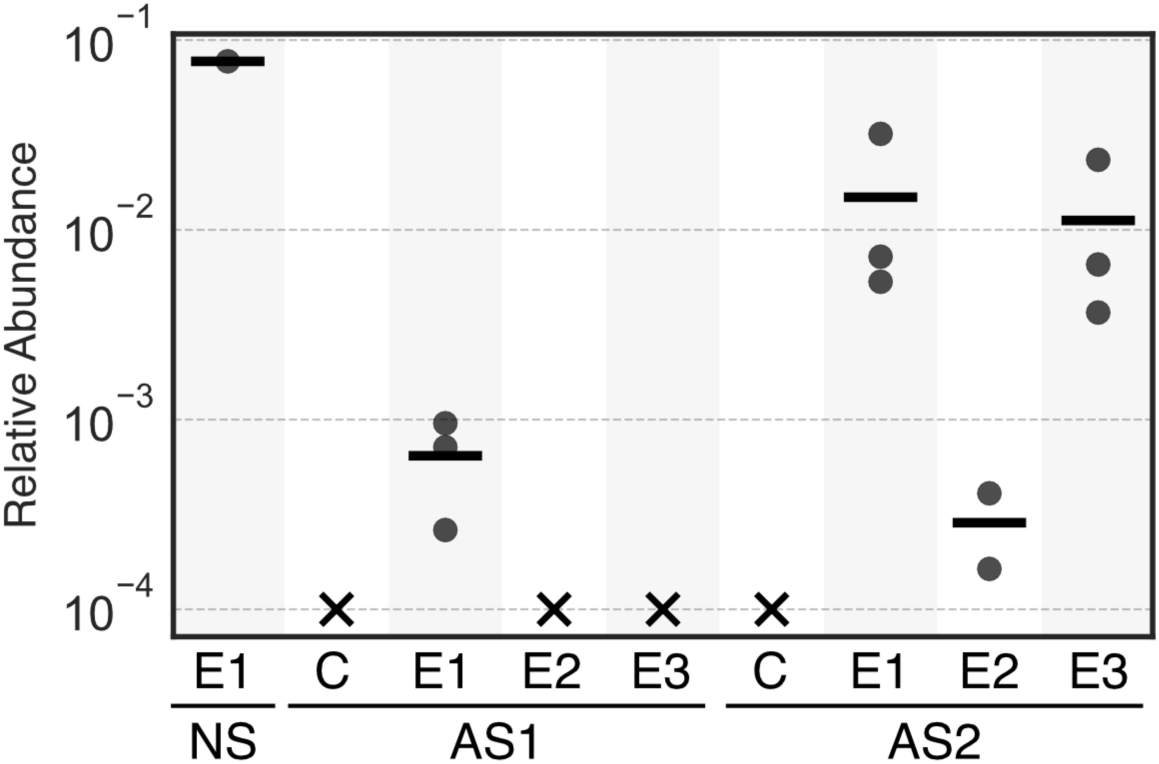
The relative abundance of *E. aureum* reads in the natural sediment (NS) as well as AS1 and AS2 that were not inoculated with *E. aureum* (C) or the primary (E1), secondary (E2), or tertiary (E3) serial enrichment of artificial sediments (n=3 for all serial enrichments except AS2-E2, n=2 for AS2-E2, n=1 for the natural sediment inoculum and the AS1 and AS2 negative controls).

To determine the microbial community diversity under the different enrichment conditions, we evaluated the alpha-diversity of each enrichment condition and beta-diversity between each enrichment condition. Species richness was represented in rarefaction curves showing that for all samples as sequencing depth increased the number of observed ASVs plateaued (**Figure S2**). At all read depths the natural sediments had higher number of ASVs than AS2 or AS1. Based on the number of observed ASVs, the species richness was highest in the natural sediment followed by AS2-E1, AS2-E3, AS2-E2, AS1-E1, AS1-E2, AS1-E3. Similar trends in species diversity were observed for both the Shannon diversity index and the Simpson diversity index (**Figure S4**). To determine how similar the microbial communities were between the natural sediment and the two artificial sediments, we assessed the beta-diversity by performing robust principal-coordinate analysis and calculating pairwise robust Aitchison distances^39^ (**Figure S5)**. AS1 sediments formed a distinct cluster while AS2 clustered closer to the natural sediment and AS1 had larger Aitchison distances relative to natural sediments. These alpha-diversity results indicate that AS2 sediments harbored a more diverse microbial community relative to AS1, and both artificial sediments represent reduced microbial community complexity relative to the natural sediment that served as the inoculum. Additionally, based on the beta-diversity the microbial community associated with AS2 was more similar to natural sediment than the AS1 community.

Looking at the most abundant genera in each sediment, we observed co-enrichment of microbes from diverse functional guilds including sulfate-reducers, iron-reducers, carbohydrate-fermenting organisms and putative nitrogen fixers (**Figure 6**). Sulfate reducers including *Desulfobulbus* and *Humidesulfovibrio* likely participate in cycling sulfate that is released by cable bacteria e-Sox and other sulfide oxidizers^40–42^. Other sulfide oxidizing bacteria including *Sulfuricurvum* are enriched along with the cable bacteria likely because these sediment conditions broadly enrich for sulfide oxidizing bacteria. The iron-reducing microbes *Pelotalea* and *Rhodoferax*, together with iron-oxidizing microbe *Leptothrix* are likely involved in Fe cycling after Fe(II) is released upon FeS oxidation^38,43^. Although *Fibrobacteraceae* possible genus 06 and *Mobilitalea* were initially present at low abundance in the natural sediment inoculum, they became stably enriched and emerged as the two most dominant genera in both AS1 and AS2. This enrichment is likely driven by the process of cellulose fermentation caused by the inclusion of the alpha-cellulose as the structuring organic substrate in the artificial sediments^44,45^. Additionally, alpha-cellulose may have also introduced additional taxa such as *Cellvibrio*, which were low abundance in the natural sediment but appeared in the negative control enrichments for both AS1 and AS2. Collectively, these results indicate that artificial sediments enrich a metabolically diverse but simplified microbiome relative to the natural sediment.

**Figure 6:**
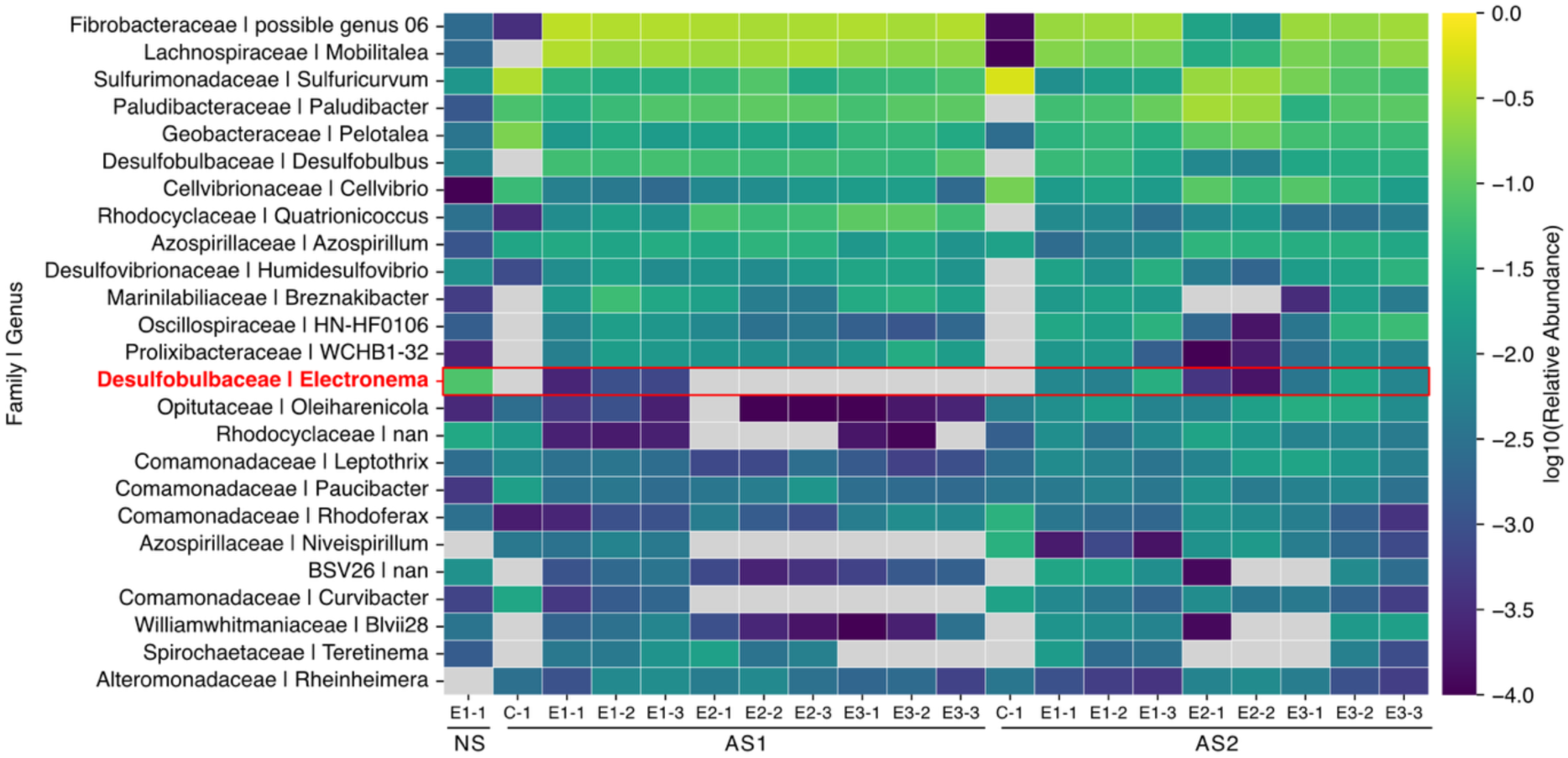
Heat map depicting the log abundance of the top 25 genera from natural sediments as well as AS1 and AS2 that were not inoculated with *E. aureum* (C) or the primary (E1), secondary (E2), or tertiary (E3) serial enrichment of artificial sediments inoculated with *E. aureum*. Data are relative abundances from individual replicate cores (n=3 for all serial enrichments except AS2-E2 which has n=2, n=1 for negative controls and the natural sediment inoculum).

Given the observation of flocking microbes in the trench slide experiments we investigated if the microbial community contained putative flocking taxa that were previously identified^38^. In total, 11 genera containing putative flocking organisms were identified in the reads for AS2 sediments including *Desulfobulbus, Humidesulfovibrio, Rhodoferax, Sideroxydans, Methylotenera, Acidovorax, Methyloversatilis, Rubrivivax, Aquabacterium, Brevundimonas,* and *Pseudopelobacter*. Many of these genera were minor members of the overall microbial community based on relative abundance (**Figure S6**). The indication of reads matching these previously identified flocking organisms corroborate the live cell imaging data. This observation illustrates how artificial sediments not only enable cultivation of cable bacteria but also recapitulate ecological interactions of the cable bacteria associated microbiome.

### Evaluating artificial sediment enrichments of marine cable bacteria

To determine if AS2 could also support e-SOx activity of marine cable bacteria, we established an enrichment of a marine strain collected from a tidal mudflat in Manasquan, NJ, USA and used this to inoculate AS2 sediments in an artificial seawater medium. See methods for details on field sampling and enrichment. We performed full-length 16S rRNA gene sequencing using nanopore long-read sequencing and extracted reads with >98% sequence identity to known cable bacteria sequences. Based on phylogeny of the 16S rRNA gene the closest related cable bacterium sequence is OBRS01813650 from a metagenomic sample^46^ sharing >98.8% sequence identity^47^, which falls into Cluster VI cable bacteria based on the taxonomy suggested by Ley *et al.*^32^ which contain the *Electrothrix* sp. (**Figure S7-8**). Additionally, to determine morphology we performed SEM on these enriched cable bacteria finding they have ∼1 μm diameter with ∼14 ridges (**Figure S9**). Based on this information, we are designating this enriched strain *Candidatus* Electrothrix sp. NJ1.

When grown in artificial sediments, *Ca*. Electrothrix sp. NJ1 behaved similarly to *E. aureum* (**Figures 7 and 1-2**). In the non-inoculated controls, the pH profile was flat over the 6 cm depth profile, while the oxygen profile decreased from 200 μM at the surface to 0 μM at a depth or ∼4 mm. In contrast, after 120 days the core inoculated with *Ca*. Electrothix sp. NJ1 showed pH profiles that are indicative of e-SOx activity with proton production below the depth of oxygen penetration and consumption above. However, in both the non-inoculated and inoculated cores the sulfide signal remained below the limit of detection. Additionally, we evaluated if AS2 could support serial enrichments of the marine cable bacterium *Ca*. Electrothrix sp. NJ1. Sediments were allowed to develop for 30 days prior to transfer to freshly prepared AS2. Following three serial transfers we assessed the geochemical profiles of all three cores at 120, 90, and 60 days post inoculation for the primary, secondary, and tertiary enrichments, respectively (**Figure 7B**). All three serial enrichments showed pH profiles indicative of e-SOx activity, while the oxygen profiles showed oxygen penetrating between 2-3 mm. In contrast to *E. aureum* enrichments, the sulfide signal remained below the limit of detection across all serial enrichments. Collectively, these results illustrate that AS2 can serve as a generalizable growth matrix for reproduceable cultivation of both freshwater and marine cable bacteria.

**Figure 7.**
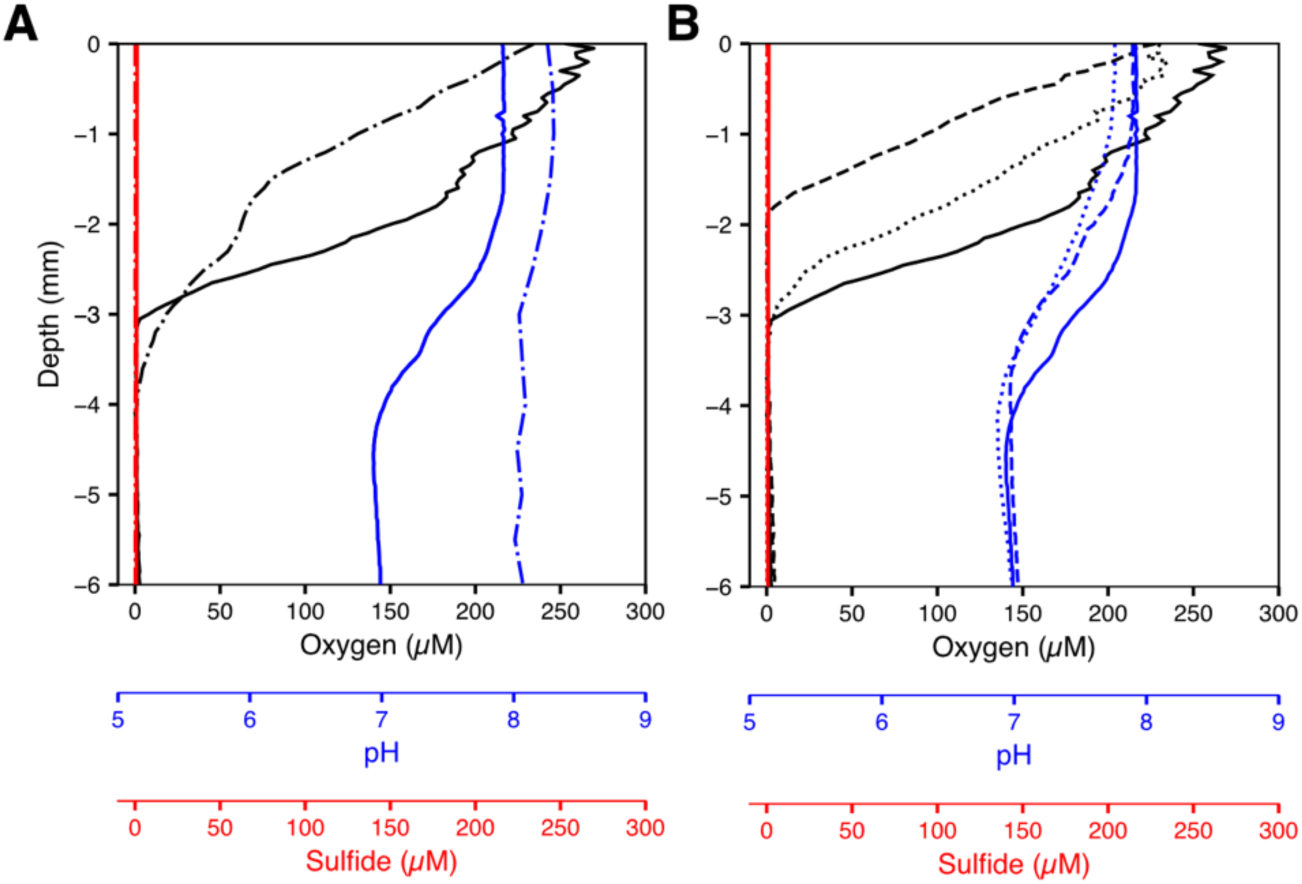
Vertical chemical profiles of artificial sediments with marine cable bacteria **(A)** AS2 inoculated with enrichments of marine *Ca*. Electrothix sp. NJ1 (solid line) or non-inoculated controls (dash-dot line). **(B)** AS2 inoculated with marine *Ca*. Electrothix sp. NJ1 with primary (solid line), secondary (dotted line), tertiary (dashed line) enrichments. AS2 sediments were submerged in aerated ASM at a salinity of 25 ppt. Measurements were taken 120, 60, and 30 days after inoculation for the primary, secondary, and tertiary enrichment, respectively. For reference, the primary enrichment is shown in both panels A and B.

## Conclusions

The artificial sediments presented herein recapitulate hallmark metabolic signatures and ecological interactions of cable bacteria that are enriched in natural sediments. The artificial sediments were shown to enable cultivation of phylogenetically diverse species of cable bacteria including both freshwater (Cluster I) and marine (Cluster VI) strains (**Figure S7-8**). In the future, artificial sediments should be evaluated if they can enable cultivation of the phylogenetically diverse cable bacteria from Clusters II-V^32^.

As shown here, artificial sediments can enable the enrichment of microbes that are not amenable to isolation using conventional culturing techniques in liquid or solid culture media. The ability to tune both the physical and chemical properties of the environment can enable the study of how the environment shapes the growth, metabolism, and ecological interactions of environmental microbes. For example, the influence of differing sulfur, nitrogen, and carbon sources on the metabolism of cable bacteria is currently underexplored due to the chemical heterogeneity of natural sediments. The artificial sediments presented here could enable direct control over the type and abundance of sulfur, nitrogen, and organic carbon compounds within the environment to interrogate cable bacteria metabolism.

Additionally, the reduced heterogeneity of artificial sediments relative to environmental samples provide controlled environments for evaluating DNA delivery approaches including conjugation and natural transformation. This could potentially enable genetic manipulation and provide a route for interrogating sequence-function relationships within cable bacteria genomes. Continued refinement of the artificial sediments may potentially lead to a route for establishing axenic cultures of cable bacteria. Together these results provide an approach for researchers to study cable bacteria and other benthic microbial communities that are recalcitrant to conventional culturing techniques in chemically defined conditions.

## Acknowledgements

We would like to thank Pia Bomholt Jensen for help with sample preparation and SEM imaging. We would like to thank Britta Poulsen for help with sample preparation and 16S rRNA gene amplicon sequencing. We would like to thank Jesper Wulff for help fabricating trench slides. We would like to thank Lillia Ryazanov for identifying the tidal flat in Manasquan, NJ. J.T.A. was partially supported by the NSF Postdoctoral Research Fellowships in Biology Program under Grant No. 2010604. This study was supported by the Office of Navy Research Grant No. N00014-24-1-2708 to J.T.A., the European Union’s Horizon Research and Innovation Program under the Marie Sklodowska-Curie grant agreement to K.A. (project 101109777, Cable Electricity O_2_ and the Danish National Research Foundation to L.P.N., A.S. and I.P.G.M. (Center for Electromicrobiology, DNRF136). ME-N acknowledges support by the WM Keck Foundation award 8626 and Gordon and Betty Moore Foundation grant 10148.

## Supplemental Information

**Figure S1.**
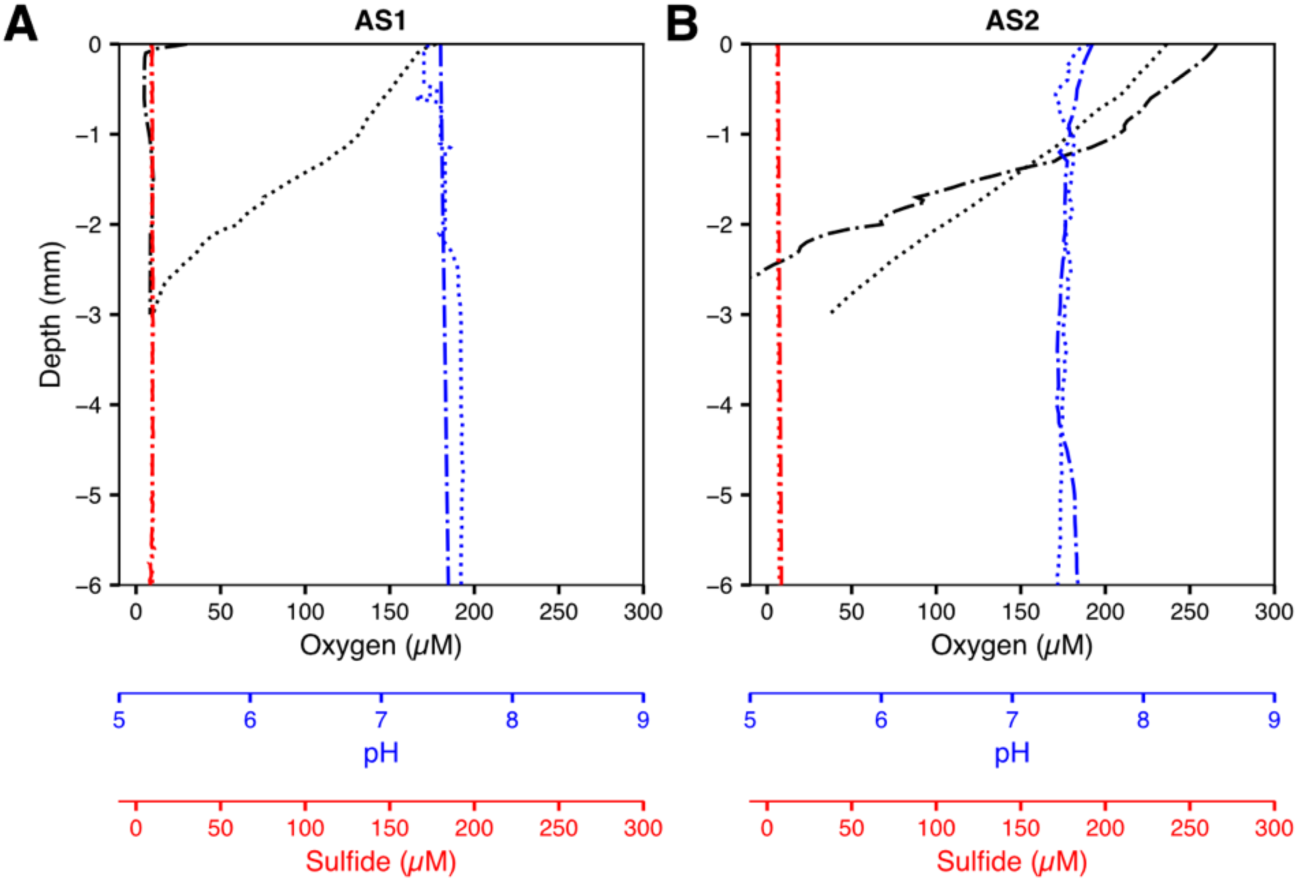
Vertical chemical profiles of non-inoculated artificial sediments. Oxygen (black), sulfide (red) and pH (blue) profiles were measured for artificial sediments for **(A)** AS1 and **(B)** AS2 of non-inoculated sediments immediately before inoculation (dotted line) and following 14 days of incubation (dashed-dot line, same data as reported in Figure 1 for reference).

**Figure S2.**
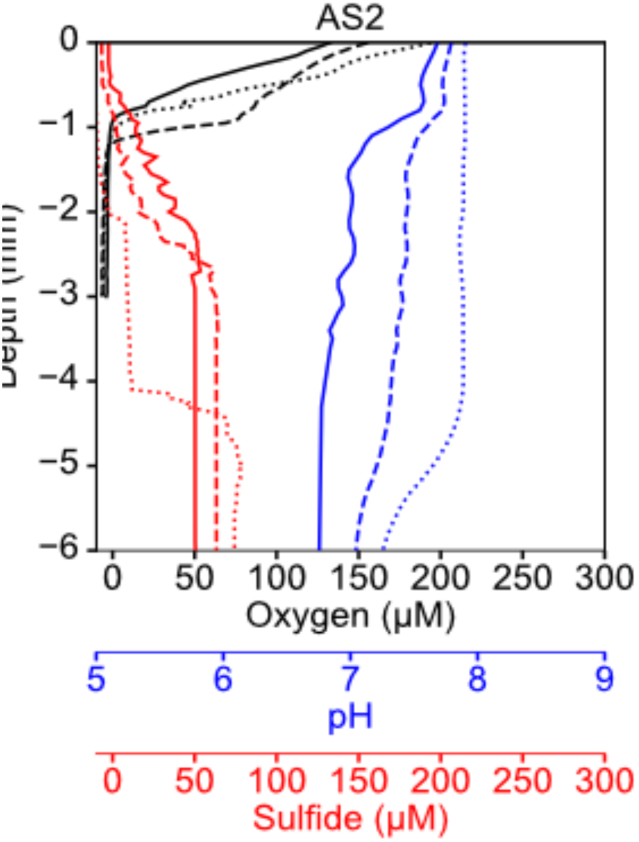
Vertical chemical profiles of artificial sediments for enrichments 4-6. Oxygen (black), sulfide (red) and pH (blue) profiles were measured for artificial sediments for AS2 of freshwater *E. aureum* for quarternary (solid line), quinary (dash-dotted line), senary (dashed line) enrichments.

**Figure S3.**
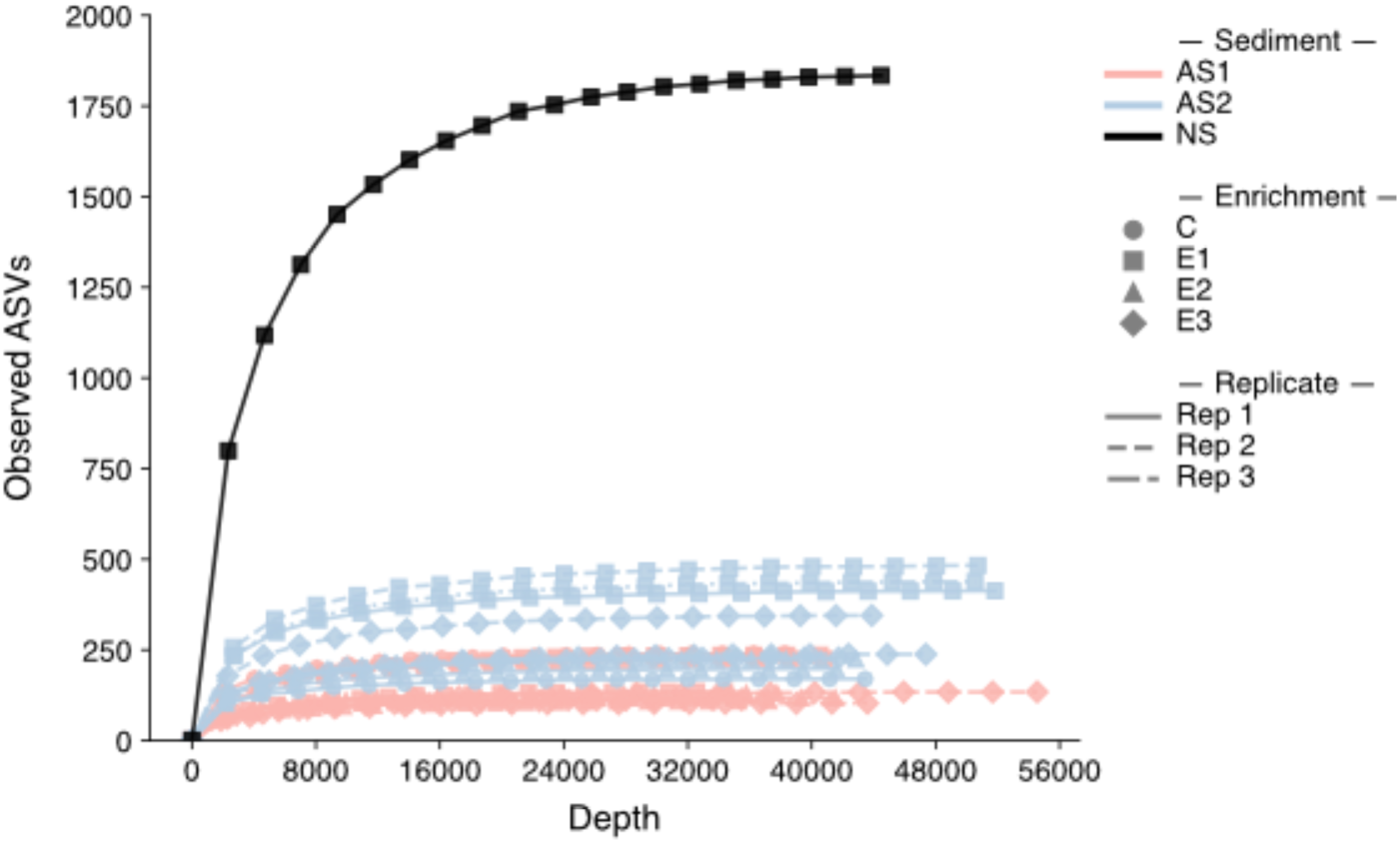
Species richness rarefaction curves. The number of observed ASVs varies as read depth increases for the natural sediment (black) and artificial sediments AS1 (red) and AS2 (blue).

**Figure S4.**
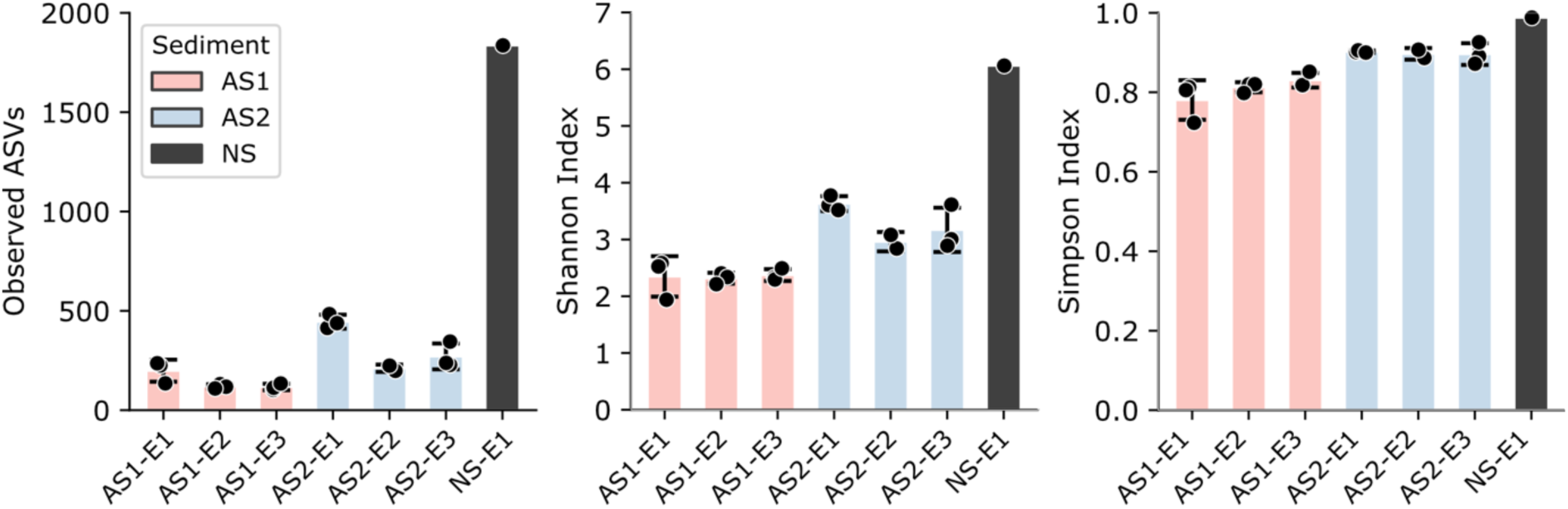
Alpha-diversity metrics (Observed ASVs, Shannon Index, and Simpson Index) for the freshwater enrichments on natural sediment (black) and artificial sediments AS1 (red) and AS2 (blue).

**Figure S5.**
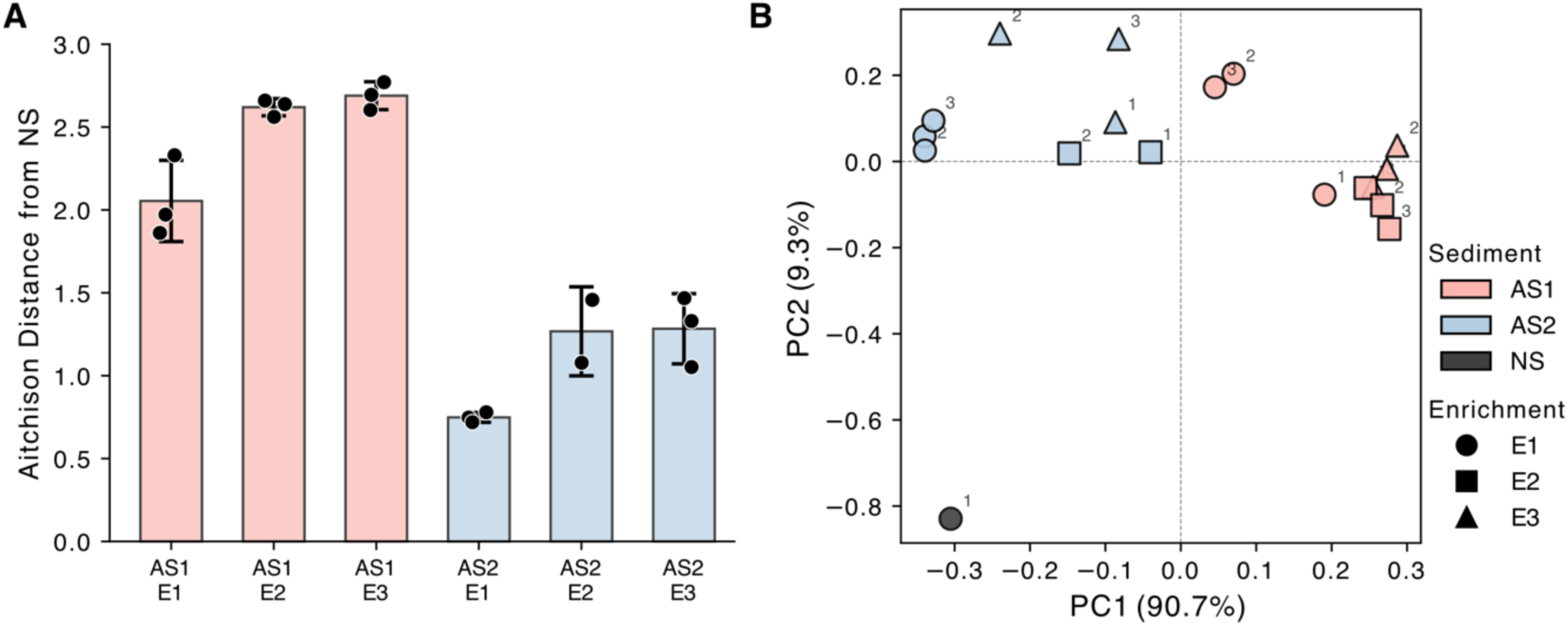
Variation in the microbial community across artificial sediment enrichments. **(A)** robust Aitchison distances^39^ between each artificial sediment enrichment relative to the natural sediment community **(B)** robust principal coordinate analysis (PC1 and PC2) among sediment enrichments. AS1 are blue, AS2 are red, NS is black. enrichment 1 are circles, enrichment 2 are squares, enrichment 3 are triangles. Bar plots and error bars represent the mean and standard deviation, respectively, and individual replicates shown as circles.

**Figure S6.**
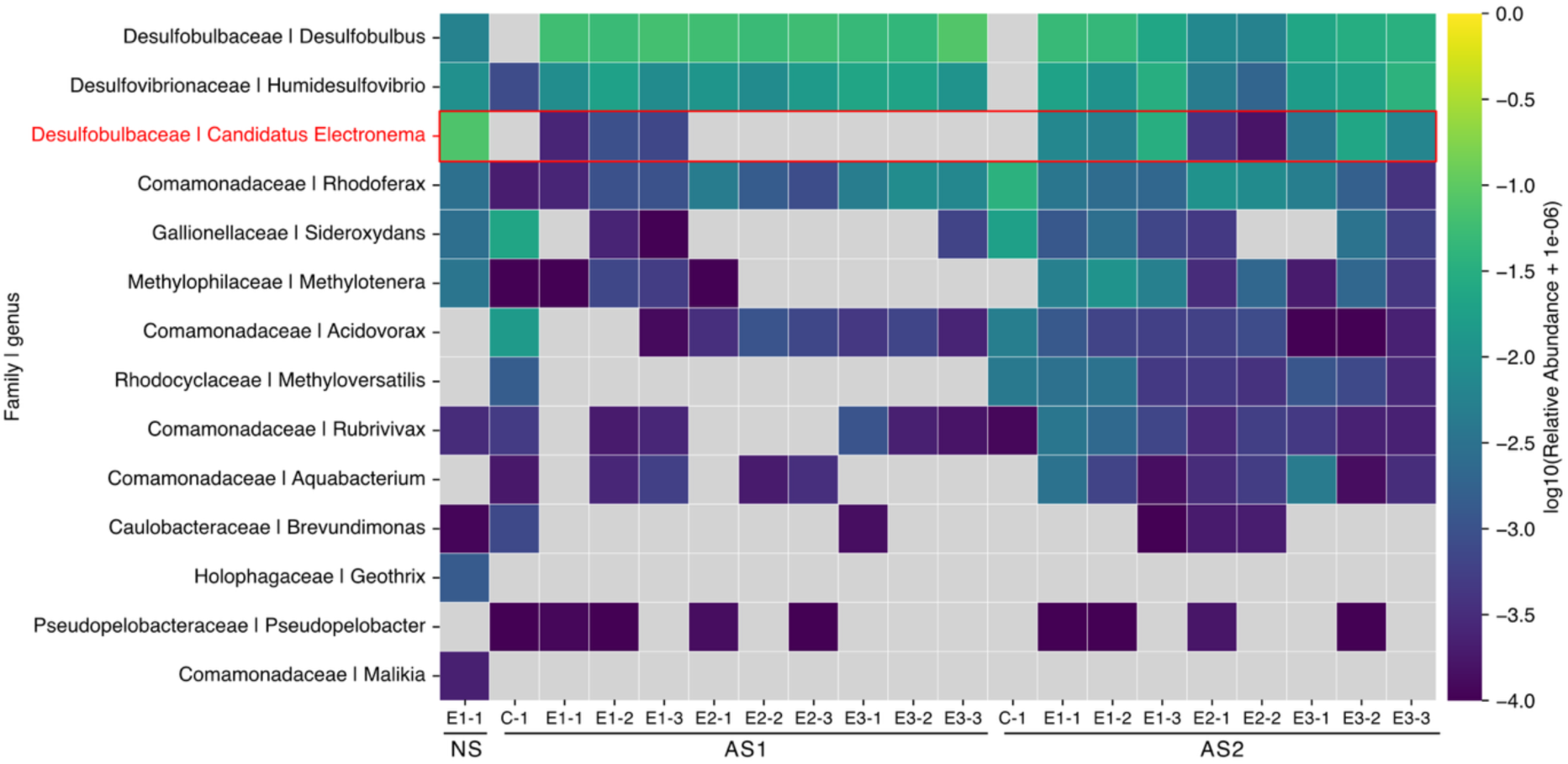
Heat map depicting the log abundance of putative flocking genera from natural sediments as well as AS1 and AS2 that were not inoculated with *E. aureum* (C) or the primary (E1), secondary (E2), or tertiary (E3) serial enrichment of artificial sediments inoculated with *E. aureum*. Data are relative abundances from individual replicate cores (n=3 for all serial enrichments except AS2-E2 which has n=2, n=1 for negative controls and the natural sediment inoculum).

**Figure S7.**
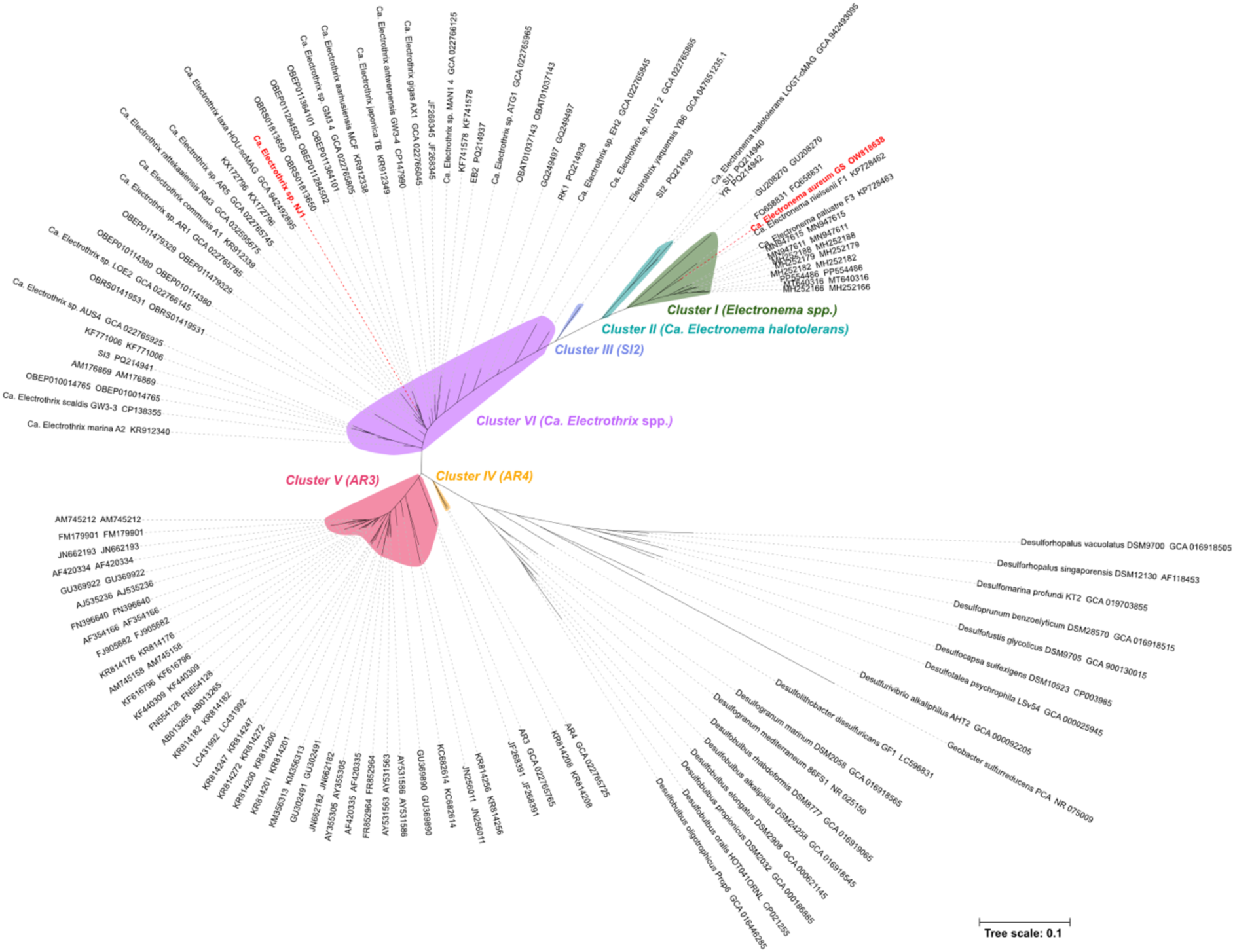
Phylogenetic tree of cable bacterial 16S rRNA genes. Phylogeny was constructed by aligning 16S rRNA gene sequences (>1200 bp) of cable bacteria and related taxa (n=107) from Ley et al.^32^ as well as the recently identified *Ca*. Electrothrix yaqonensis YB6^33^ and the *Ca*. Electrothrix sp. NJ1 recovered in this study using MUSCLE^31^. A maximum likelihood tree was constructed with IQ-TREE (v1.6.12)^34^, employing the TIM3e+I+G4 model and 1,000 ultrafast bootstrap replicates. Tree was visualized using iTOL^35^. The two cable bacteria used in this study *Electronema aureum* GS and *Ca.* Electrothrix sp. NJ1 are colored in red. Scale bar represents 10% estimated sequence divergence.

**Figure S8.**
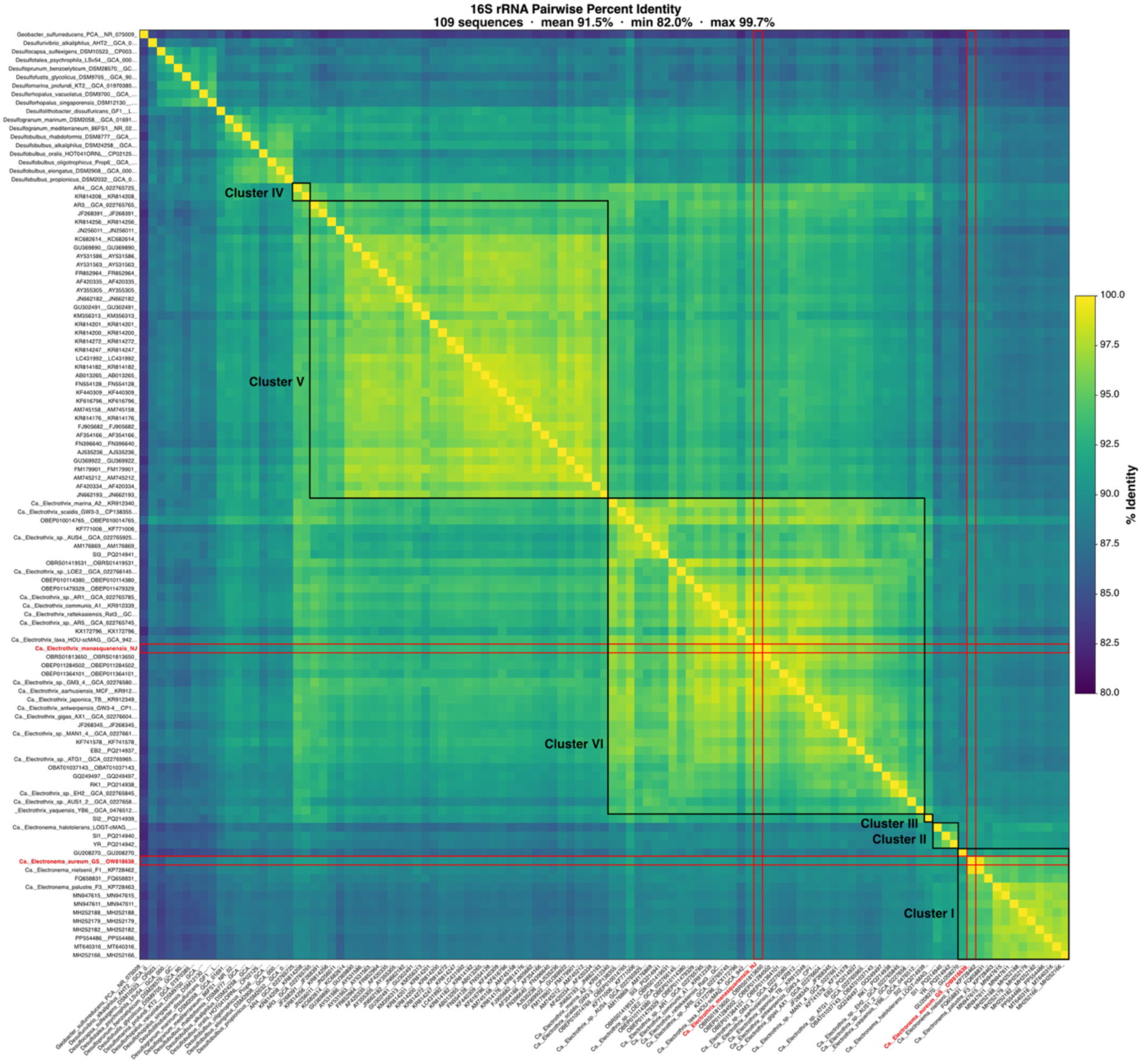
Pairwise sequence identity matrix of cable bacteria 16s rRNA genes. Clustering is based on the phylogenetic tree in Figure S5. The two cable bacteria used in this study *Electronema aureum* and *Ca.* Electrothrix sp. NJ1 are colored in red.

**Figure S9.**
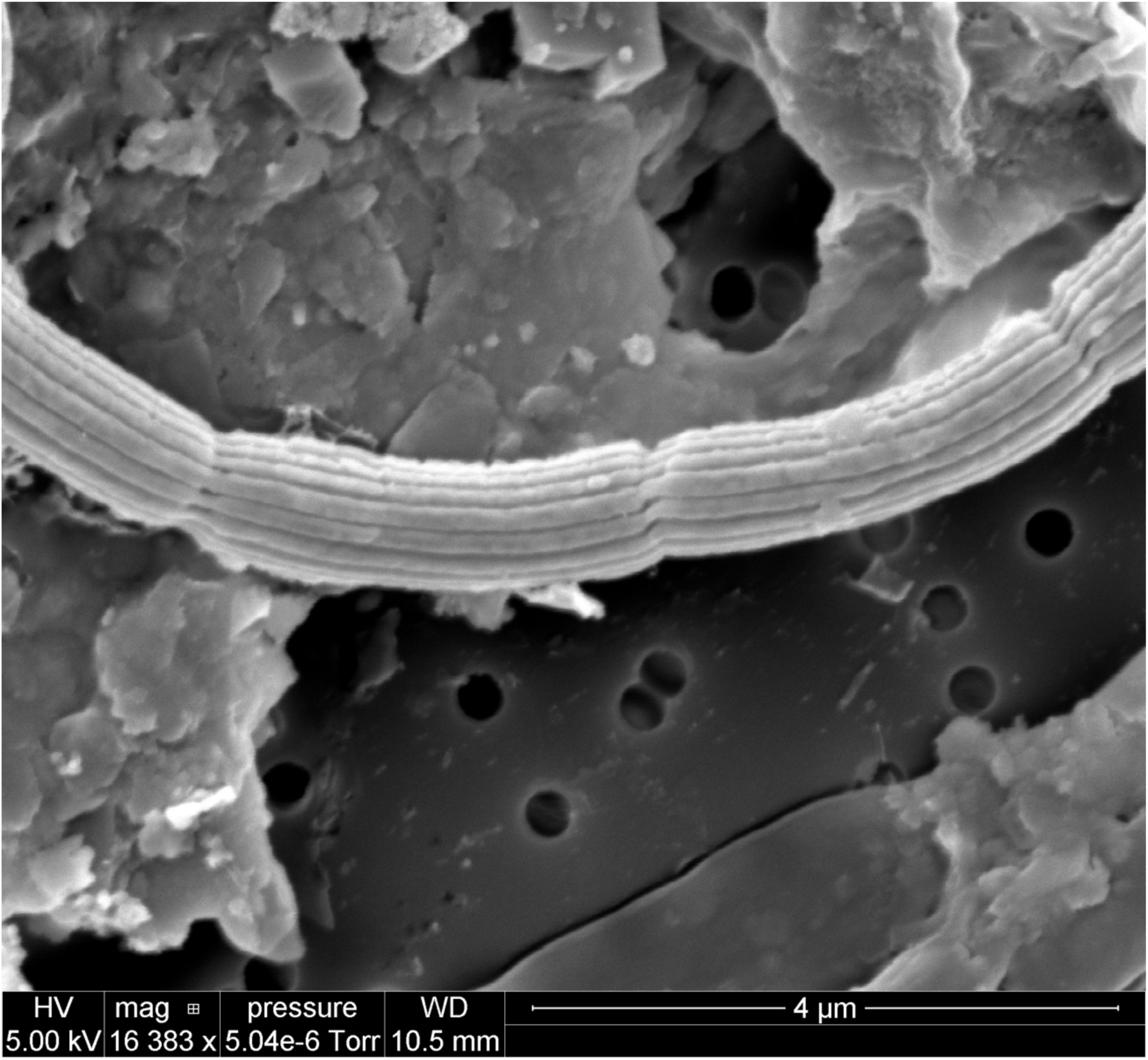
Scanning electron micrographs of *Ca.* Electrothrix sp. NJ1 enriched in natural sediments.

**Video S1.**
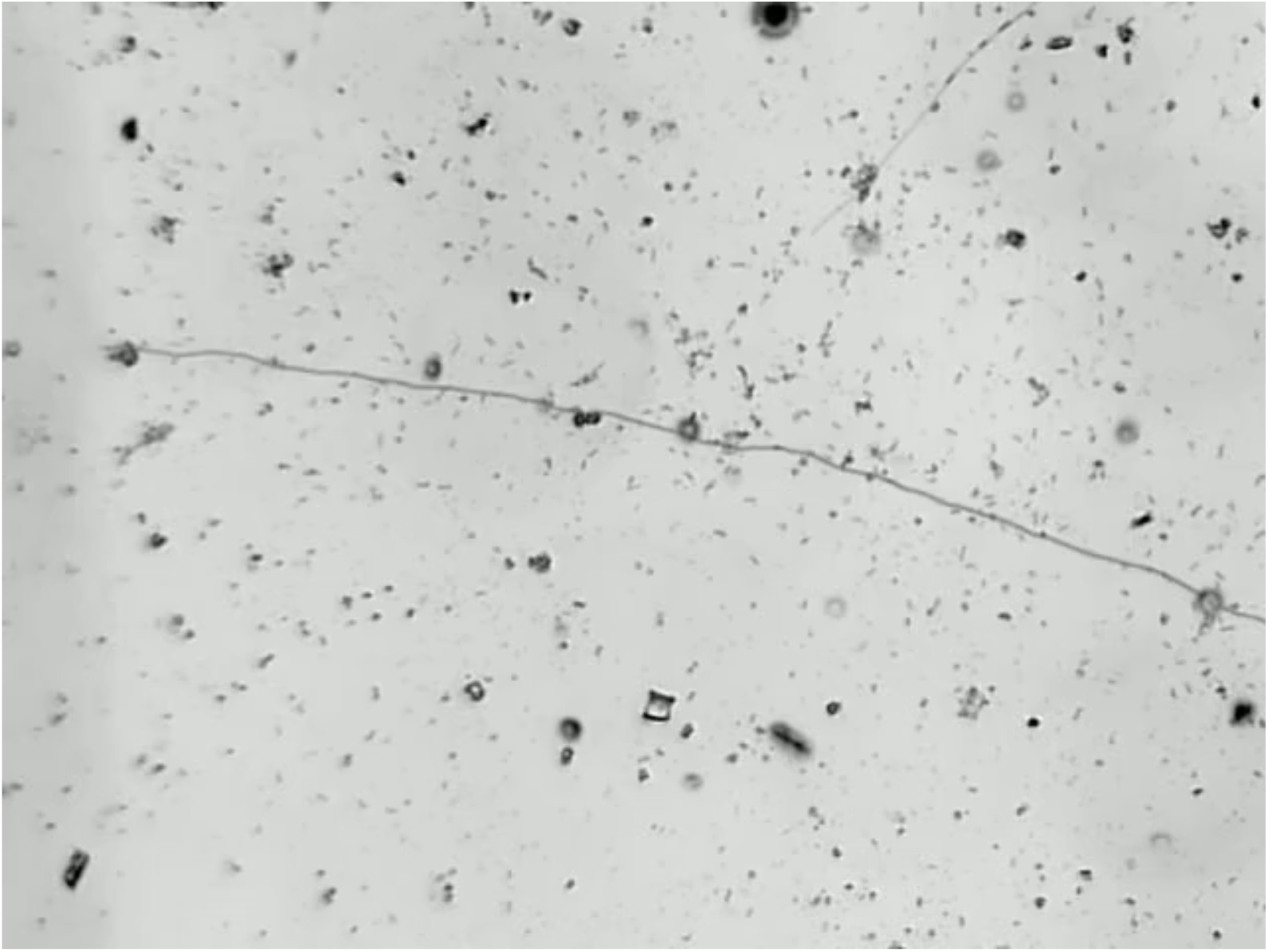
Live cell phase contrast microscopy of single celled organisms flocking around an *E. aureum* filament grown in AS2 in a trench slide device. Video is at 8x speed.

**Table S1.**
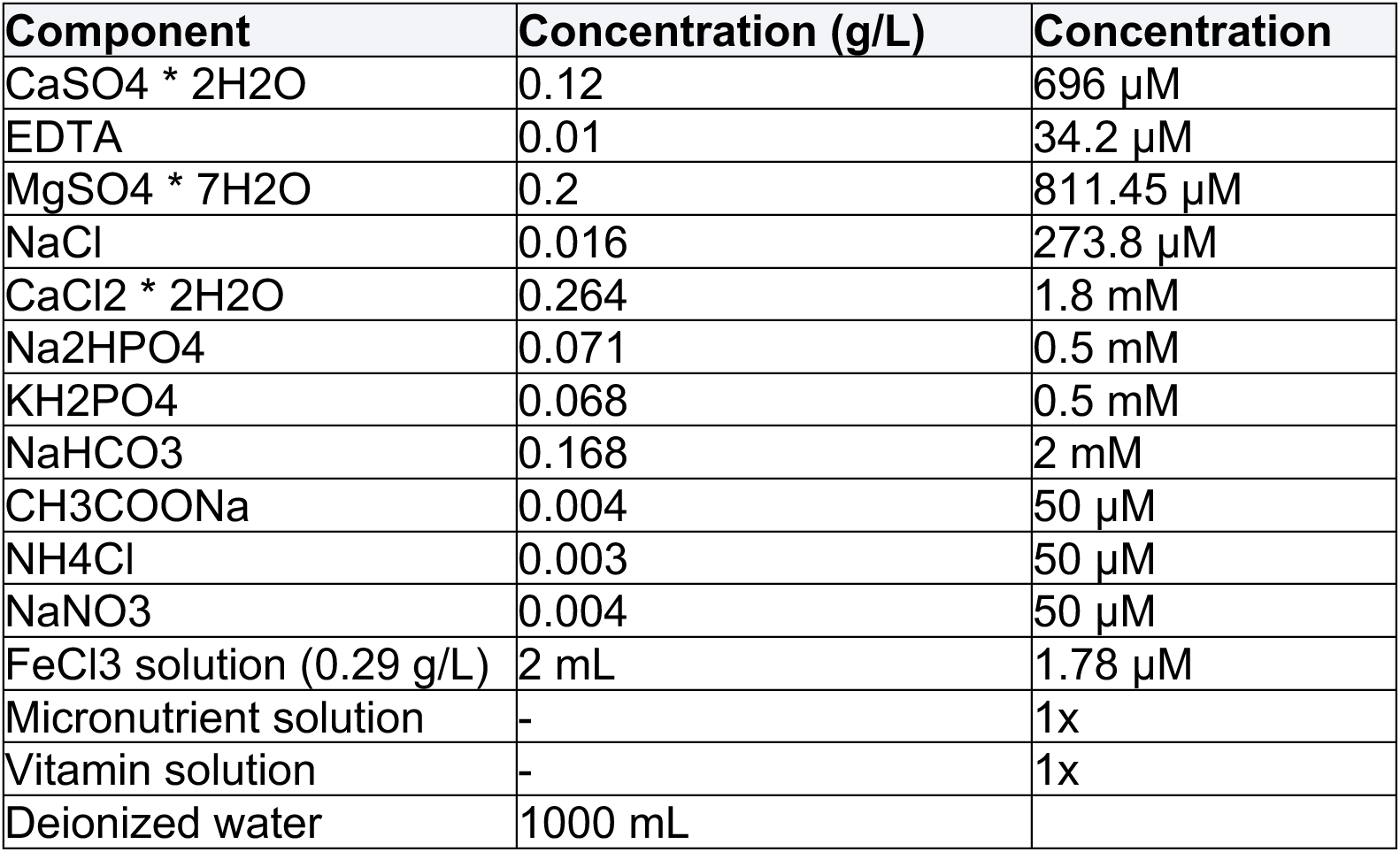
Freshwater benthic medium (FWBM) recipe.

**Table S2.** Micronutrient solution recipe. The 1000x micronutrient solution consisted of the following (per 1000 mL): sulfuric acid (0.5 mL), manganese sulfate monohydrate (2.2 g), zinc sulfate heptahydrate (0.5 g), boric acid (0.5 g), cupric sulfate pentahydrate (0.025 g), sodium molybdate dihydrate (0.025 g), cobalt chloride hexahydrate (0.045 g), nickel chloride hexahydrate (0.025 g). The solution was filter sterilized and stored at 4°C until use.

| Component | Concentration (mL/L or g/L) |
| --- | --- |
| H<sub>2</sub>SO<sub>4</sub>, sulfuric acid (>98%) | 0.5 mL |
| MnSO<sub>4</sub> * H<sub>2</sub>O | 2.2 g |
| ZnSO<sub>4</sub> * 7 H<sub>2</sub>O | 0.5 g |
| H<sub>3</sub>BO<sub>3</sub> | 0.5 g |
| CuSO<sub>4</sub> * 5 H<sub>2</sub>O | 0.025 g |
| Na<sub>2</sub>MoO<sub>4</sub> * 2 H<sub>2</sub>O | 0.025 g |
| CoCl<sub>2</sub> * 6 H<sub>2</sub>O | 0.045 g |
| NiCl<sub>2</sub>* 6 H<sub>2</sub>O | 0.025 g |
| deionized water | 1000 mL |

**Table S3.**
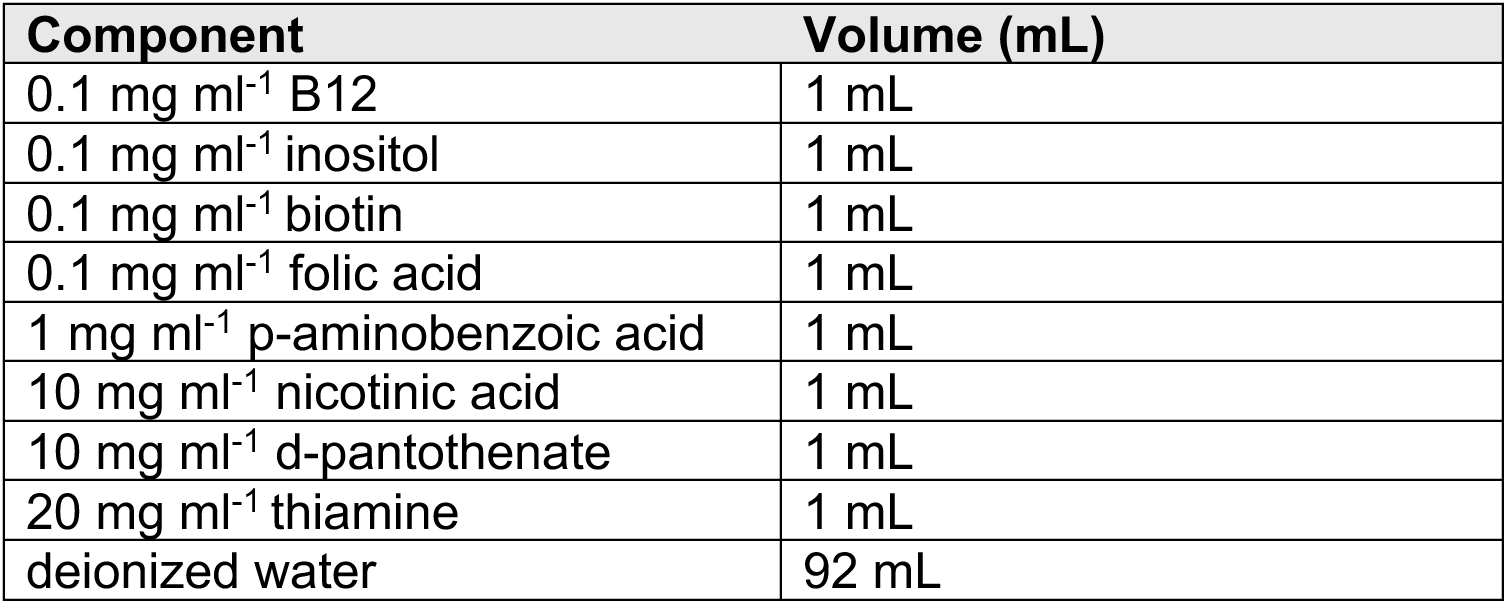
Vitamin solution recipe. The 2400x vitamin solution consisted of individual stock solutions of the following (per 100 mL): 0.1 mg ml^−1^ B12, 0.1 mg ml^−1^ inositol, 0.1 mg ml^−1^ biotin, 0.1 mg ml^−1^ folic acid, 1 mg ml^−1^ p-aminobenzoic acid, 10 mg ml^−1^ nicotinic acid, 10 mg ml^−1^ d-pantothenate, 20 mg ml^−1^ thiamine. Stock solutions of individual components were initially prepared in deionized water and stored at −80°C until use. To obtain the final stock solution, 1 ml each of the above stock solutions were added to 92 ml deionized water, followed by filter sterilization. The solution was stored at 4°C until use.

| Component | Volume (mL) |
| --- | --- |
| 0.1 mg ml<sup>-1</sup> B12 | 1 mL |
| 0.1 mg ml<sup>-1</sup> inositol | 1 mL |
| 0.1 mg ml<sup>-1</sup> biotin | 1 mL |
| 0.1 mg ml<sup>-1</sup> folic acid | 1 mL |
| 1 mg ml<sup>-1</sup> p-aminobenzoic acid | 1 mL |
| 10 mg ml<sup>-1</sup> nicotinic acid | 1 mL |
| 10 mg ml<sup>-1</sup> d-pantothenate | 1 mL |
| 20 mg ml<sup>-1</sup> thiamine | 1 mL |
| deionized water | 92 mL |

